# Treatment response prediction: Is model selection unreliable?

**DOI:** 10.1101/2022.03.19.483454

**Authors:** David Augustin, Ken Wang, Antje-Christine Walz, Ben Lambert, Michael Clerx, Martin Robinson, David Gavaghan

**Affiliations:** Computer Science, University of Oxford, Parks Road, Oxford, OX1 3QD, UK; Research and Early Development, F. Hoffmann-La Roche AG, Grenzacherstrasse 124, Basel, 4070, Switzerland; Faculty of Science, University of Nottingham, University Park, Nottingham, NG7 2RD, UK

**Keywords:** Probabilistic Model Averaging, Model Selection, Model Averaging, PKPD Modelling, Dose Response, Dose Selection, Nonlinear Dynamics

## Abstract

Quantitative modelling has become an essential part of the drug development pipeline. In particular, pharmacokinetic and pharmacodynamic models are used to predict treatment responses in order to optimise clinical trials and assess the safety and efficacy of dosing regimens across patients. It is therefore crucial that treatment response predictions are reliable. However, the data available to fit models are often limited, which can leave considerable uncertainty about the best model to use. Common practice is to select the model that is most consistent with the observed data based on the Akaike information criterion (AIC). Another popular approach is to average the predictions across the subset of models consistent with the data. In this article, we argue that both approaches can lead to unreliable predictions, as treatment responses typically display nonlinear dynamics, so models can be consistent with the observed dynamics, whilst predicting incorrect treatment responses. This is especially the case when predicting treatment responses for either times or dosing regimens that go beyond the observed dynamics. Across a range of experiments on both real laboratory data and synthetically derived data on *Neisseria gonorrhoeae* response to ciprofloxacin, we show that probabilistic averaging of models results in more reliable treatment response predictions.

## 1 Introduction

Modelling of pharmacological processes has become an essential tool in the pharmaceutical drug development pipeline [1–3]. In the early stages of development, models are, for example, used to optimise clinical trials by predicting clinical efficacy and safety of dosing regimens [4]. In later stages, models help to quantify the inter-individual treatment variability across patients and assess the need for dose adjustments in specific populations [5, 6]. More recently, it has also been suggested that models could be used to recommend personalised dosing strategies in clinical practice [7, 8].

While the potential of modelling for accelerated drug development and better treatment of patients has been demonstrated [9], the availability of time-resolved preclinical or clinical data, needed for the training of models, is often limited. This impairs our ability to use models to learn and predict the nonlinear treatment response dynamics of patients. To overcome the shortage of data, mechanistic PKPD models incorporate biological and pharmacological mechanisms informed by previous experiments [10–13]. These mechanistic models are able to learn treatment response dynamics more efficiently from limited data.

However, frequently it is not clear which pharmacological processes are the most relevant for assessing the treatment response of patients and, thus, multiple candidate models exist [14]. In these situations, we can be left with considerable uncertainty about the correct model choice, which can have far-reaching consequences for the reliability of model predictions [15]. Common practice is to select the model for predictions that is the most consistent with the available data based on the AIC score [16]. More recently, it has also been proposed to average the predictions of candidate models according to their AIC scores. Uster et al. introduce, for example, the model averaging algorithm (MAA) where treatment responses are predicted based on the weighted average of the mean predictions of candidate models [17]. In contrast, other authors have proposed to average the full treatment response distributions [18]. We will refer to the latter as probabilistic averaging of models (PAM), and describe both MAA and PAM in more detail in Sections 2.3 and 2.4.

In this article, we argue that optimal consistency with the observed treatment response does not imply that a model extrapolates well to the future or to unobserved dosing regimens. In particular, we demonstrate that model selection frequently fails to identify the correct candidate model when multiple models are close approximations of the observed treatment response. We also show that MAA only improves over model selection in special cases, where all candidate models with substantial weights reduce to the qualitatively similar dynamics. For more reliable treatment response predictions, we propose the use of the PAM framework as this approach is the only one that reflects the uncertainty of treatment response predictions. We show that PAM outperforms model selection and MAA in the analysis of *in vitro* time kill data, where cultures of *N. gonorrhoeae* are exposed to ciprofloxacin. In a subsequent synthetic study we analyse this behaviour in more depth.

## 2 Methods

In this section, we introduce PAM, after describing the standard PKPD modelling framework and common model selection/averaging strategies. Subsequently, in Sections 2.6-2.7, we present a typical PKPD modelling example — the treatment of bacterial infections — which we use to demonstrate the predictive performance of PAM and other model selection/averaging strategies.

### 2.1 Modelling framework

Pharmacokinetic and pharmacodynamic (PKPD) models are (semi-)mechanistic or empirical descriptions of dose-exposure-response dynamics in terms of ordinary differential and algebraic equations

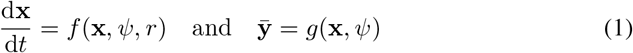

with initial conditions **x**(*t* = 0) = **x**_0_ [19–21]. **x** denotes the dynamical states, *ψ* the model parameters, *r* the dosing regimen and *t* the time.^1^ Typical dynamical states in PKPD models include the drug amount in the blood or other tissues and biomarkers for the disease progression. Common model parameters are biological, physiological or drug-specific properties. In general, not all dynamical states **x** can be observed directly (or at all). We therefore denote the measurable quantities of the treatment response by 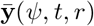, whose time-course are determined by the dynamical states through the mapping *g*. For example, in the bacterial infection use case in Section 2.5, only the total count of the bacterial population can be measured, whereas **x** may model the bacterial count of specific subpopulations. While Equation 1 may be of semi-mechanistic or empirical nature, we will refer to it as the *mechanistic model*.

In order to estimate the parameters *ψ* from *n* observations of the treatment response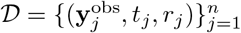, it is customary to model the discrepancy between the mechanistic model predictions and the observations with an error model. This error model converts the PKPD model to a probabilistic model

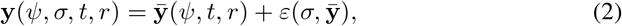

where *ε* models the random deviations of the measurable values **y** from the mechanistic model predictions 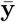. The observed values **y**^obs^ are assumed to be realisations of the measurable values **y**. The randomness of the measurable values with respect to the mechanistic model predictions reflects measurement noise and possible misspecifications of the mechanistic model [22, 23]. A common error model choice is, for example, to assume that the discrepancies *ε* are i.i.d. Gaussian distributed. Equation 2 defines a distribution *p*(**y**|*ψ, σ, t, r*) of measurable values, whose expectation is equal to the predictions of the mechanistic model 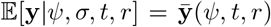.^2^ *σ* denotes the additional parameters of the error model.^3^

From this probabilistic model, maximum likelihood estimates (MLEs) of the model parameters 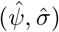 can be derived by maximising their log-likelihood

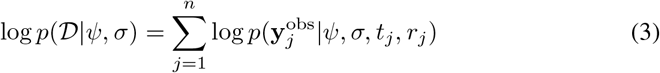

with respect to the data *𝒟*.^4^ These MLEs are often used for prediction, e.g. using the mechanistic model predictions, 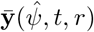, or the probabilistic model distributions, 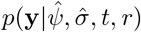 [11, 12, 24].

However, the data *𝒟* leaves uncertainty about the model parameter estimates, which is unaccounted for by the MLEs. More reliable predictions can be obtained by estimating the effect of this *parametric uncertainty* for the model predictions [25]. The parametric uncertainty can, for example, be quantified by profiling the likelihood surface or by using Bayesian inference techniques [25, 26]. Regardless of the approach, the uncertainty of the parameters can be expressed in terms of a distribution of parameter values *p*(*ψ, σ*|*𝒟*).^5^ Averaging the treatment response predictions over all parameter values results in more reliable predictions

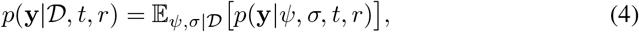

where 𝔼_*ψ,σ*|*𝒟*_ [·] denotes the expectation with respect to the parameter distribution *p*(*ψ, σ*|*𝒟*). In this article, we use Bayesian inference to estimate the distribution of parameter values in terms of the posterior parameter distribution

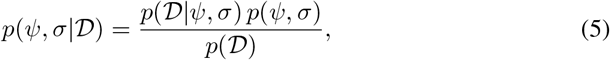

where *p*(*ψ, σ*) is the prior distribution of the model parameters and quantifies the knowledge about possible values before fitting the model to data. *p*(*𝒟*) is the model evidence and serves as a normalisation factor of the posterior distribution [27].

The model selection/averaging strategies introduced in the next sections do not require the estimation of parametric uncertainty, and the reader may replace *p*(**y**|*𝒟, t, r*) by the maximum likelihood distribution 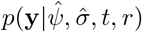 in any of the following expressions.

### 2.2 Model selection

If multiple candidate models for the treatment response dynamics exist, a widely used modelling strategy is model selection (MS) based on the Akaike information criterion (AIC) score

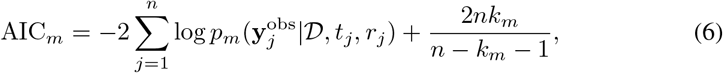

where *k*_*m*_ is the total number of model parameters [28, 29]. The subscript *m* labels the candidate models, i.e. the choice of the mechanistic model-error model pair.^6^ The AIC score estimates the cross entropy between the data-generating process and the candidate model (up to a constant factor), and can therefore be used to quantify which model captures the most information about the observed treatment response dynamics. The model with the lowest AIC score is estimated to be the most informative and is for that reason commonly selected for future predictions, e.g. in [16].

### 2.3 The model averaging algorithm

Similar to parametric uncertainty, the data *𝒟* can leave considerable uncertainty about the best candidate model choice, which is unaccounted for by MS [15]. An alternative approach that recognises this *structural uncertainty* about the candidate models is the model averaging algorithm (MAA) introduced in [17]. In this approach, the means of the candidate models are weighted by their relative likelihood, *w*_*m*_, and averaged to predict treatment responses [30]

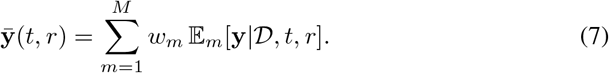

Here, 𝔼_*m*_[**y**|*𝒟, t, r*] denotes the expectation of **y** with respect to the candidate model *p*_*m*_(**y**|*𝒟, t, r*). The relative likelihoods of the models are estimated using a partition function-like normalisation of the AIC scores [31]

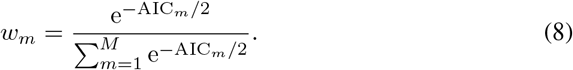

Note that for the maximum likelihood approach the expectation of **y** is equal to the mechanistic model predictions, 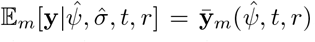, such that Equation 7 reduces to the weighted average of the mechanistic model outputs, 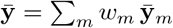.

### 2.4 Probabilistic averaging of models

MAA incorporates structural uncertainty by taking a weighted average of the candidate models’ mean predictions (see Equation 7), but it provides no overall uncertainty quantification (see Figure 1). Probabilistic averaging of models (PAM) is a model averaging strategy that is able to capture this predictive uncertainty. PAM was introduced by Akaike in 1978 and is also known as ‘model averaging’ [31, 32]. However, since different notions of model averaging have been established in the literature [17, 18, 33–35], we refer to the averaging of predictive distributions as PAM.

**Figure 1:**
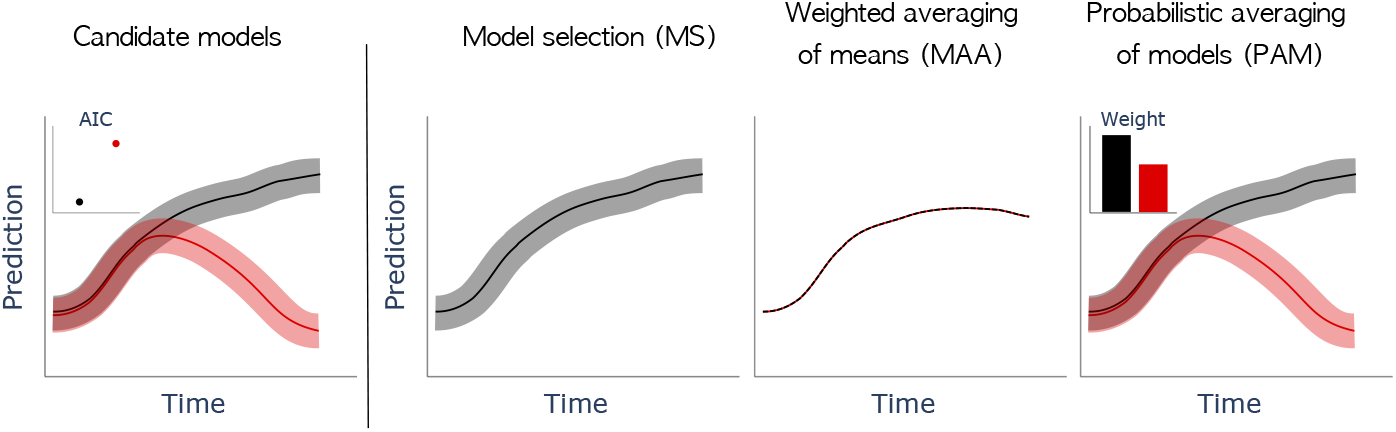
Schematic comparison of model selection/averaging methods. The left panel shows predictions of two candidate models and their AIC scores after learning from data. The next panel shows the MS predictions where the model with the lower AIC score (here the black model) is selected. The third panel shows the MAA predictions where the individual mean predictions are weighted by the model likelihoods/weights and averaged. The last panel illustrates the proposed PAM approach. In all panels the solid line represents the mean prediction of the model and the shaded areas the 5th to 95th percentiles (except for MAA which does not estimate the predictive uncertainty).

Similar to MS, PAM disregards candidate models that are inconsistent with the observed data. However, as opposed to only choosing the model with the lowest AIC score (highest likelihood), PAM bases its predictions on all models which have a substantial likelihood. To this end, PAM estimates the relative model likelihoods identically to MAA, see Equation 8. An important difference to MAA is, however, that PAM averages the probability densities, and not the mean predictions of the models

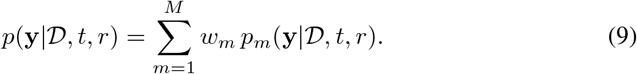

As a result, PAM is able to quantify the uncertainty of the treatment response predictions. This form of distribution average also appears in other model averaging frameworks, such as Bayesian model averaging (BMA). In BMA the cross entropy-based model weights are replaced by estimates derived from the model evidence [15, 36]. However, recently Fong and Holmes showed that the log-model evidence can be interpreted as a Bayesian approximation of the cross entropy [37], which motivates PAM not just from an information theoretic perspective, but also from Bayesian inference. Note that the mean prediction of PAM is equal to the MAA prediction, which is why MAA can be seen as a special case of PAM.

In practice, the time evolution of *p*(**y**|*𝒟, t, r*) can be challenging to visualise, which is why we illustrate the predictions of the individual candidate models together with their relative likelihoods instead (see e.g. right panel Figure 1). When the relative likelihood of a candidate model is below a threshold of *w*_thresh_ = 10^−3^, we omit the model’s prediction from the plot.

### 2.5 Candidate models

To demonstrate the consequences of MS, MAA and PAM for the reliability of treatment response predictions, we consider a typical PKPD modelling problem — the treatment of bacterial infections. Typical responses of bacterial populations to antimicrobials include an increased death rate (K), the temporary transition into a non-proliferating/persistent state (P), and the development of resistance to the treatment (R). These response mechanisms can be combined to formulate candidate models for the observed treatment response (K, KP, KR; see Figure 2) [24].

**Figure 2:**
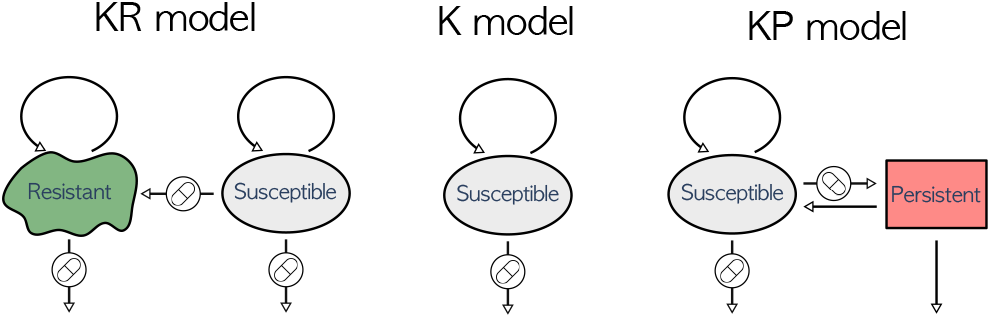
Candidate models for bacterial growth under drug exposure. Arrows tagged with a pill indicate drug-induced transitions, while other transitions are drug-independent. Self-loops represent growth, and arrows pointing downwards indicate death.

In principle, all response mechanisms may occur simultaneously, but our data (see Section 2.6) are not sufficient to identify the parameters of all mechanisms at the same time. We nevertheless define a KPR model and view the candidate models as special cases. We define the KPR model as

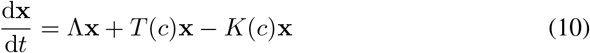

with **x** = (*x*_*s*_, *x*_*p*_, *x*_*r*_)^*t*^, where *x*_*s*_, *x*_*p*_ and *x*_*r*_ denote the counts of the susceptible (*s*), persistent (*p*) and resistant (*r*) bacteria, respectively. Experimentally, the subpopulations are not distinguished, which is why we define the measurable bacterial count as the sum of the individual counts 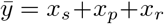. Λ = diag(*λ*_*s*_, −*λ*_*p*_, *λ*_*r*_) describes the baseline net-growth of the subpopulations, and *K* = diag(*κ*_*s*_(*c*), 0, *κ*_*r*_(*c*)) captures the drug-induced death rates. The transition rates between the subpopulations are defined as

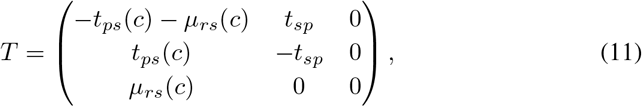

where *µ*_*rs*_ denotes the mutation rate from *s* to *r*, and *t*_*sp*_ and *t*_*ps*_ are the transition rates from *p* to *s* and *s* to *p*, respectively. The initial counts of the bacterial subpopulations are denoted by **x**_0_ = (*x*_*s*,0_, *x*_*p*,0_, *x*_*r*,0_)^*t*^. We refer to the collection of all parameters as *ψ*. From this KPR model, we can derive the K, KP and KR models according to Table 1. The dependence of the rates (*µ*_*rs*_, *t*_*ps*_, *κ*_*s*_, *κ*_*r*_) on the concentration *c* is assumed to be sigmoidal. However, for some rates the *in vitro* data does not identify both the maximal kill rate and the half maximal effect concentration. Those rates are assumed to depend linearly on the drug concentration (see Table 1). Our definition of the candidate models are minor adaptations of previously introduced models from Nielsen and Friberg [24].

**Table 1.**
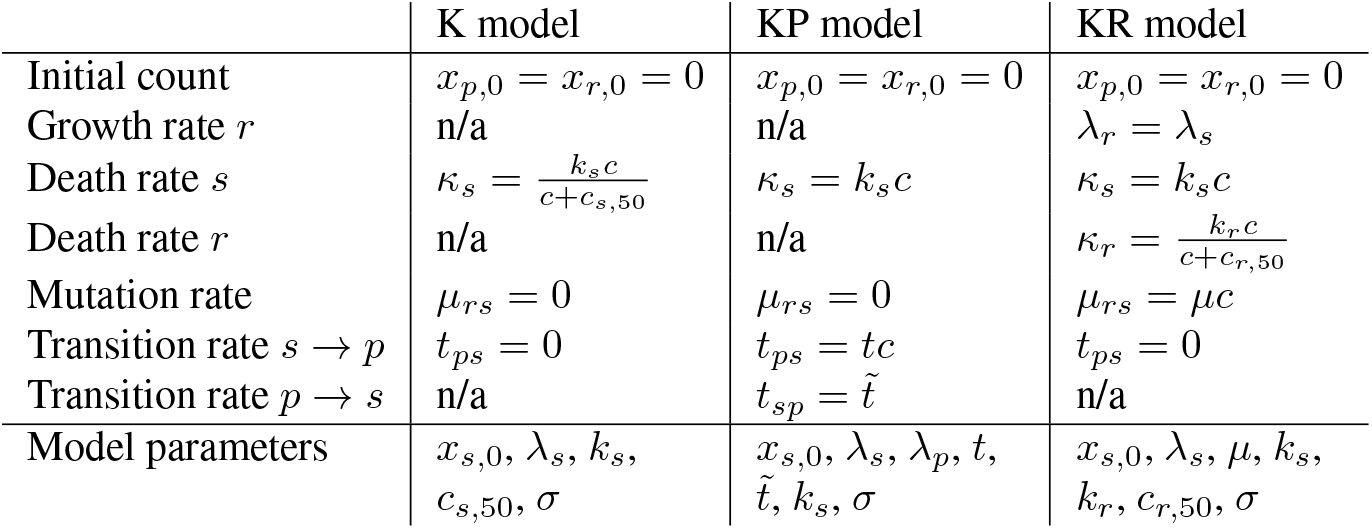
Definitions of candidate models as special cases of the KPR model in Equations 10 and 11. All model parameters are non-negative constants that need to be estimated from data.

|  | K model | KP model | KR model |
| --- | --- | --- | --- |
| Initial count | $x_{p,0} = x_{r,0} = 0$ | $x_{p,0} = x_{r,0} = 0$ | $x_{p,0} = x_{r,0} = 0$ |
| Growth rate $r$ | n/a | n/a | $\lambda_r = \lambda_s$ |
| Death rate $s$ | $\kappa_s = \frac{k_s c}{c + c_{s,50}}$ | $\kappa_s = k_s c$ | $\kappa_s = k_s c$ |
| Death rate $r$ | n/a | n/a | $\kappa_r = \frac{k_r c}{c + c_{r,50}}$ |
| Mutation rate | $\mu_{rs} = 0$ | $\mu_{rs} = 0$ | $\mu_{rs} = \mu c$ |
| Transition rate $s \rightarrow p$ | $t_{ps} = 0$ | $t_{ps} = t c$ | $t_{ps} = 0$ |
| Transition rate $p \rightarrow s$ | n/a | $t_{sp} = \tilde{t}$ | n/a |
| Model parameters | $x_{s,0}, \lambda_s, k_s,$<br>$c_{s,50}, \sigma$ | $x_{s,0}, \lambda_s, \lambda_p, t,$<br>$\tilde{t}, k_s, \sigma$ | $x_{s,0}, \lambda_s, \mu, k_s,$<br>$k_r, c_{r,50}, \sigma$ |

We choose to model the error of the mechanistic model for all candidate models with a lognormal distribution.^7^ As a result, the distribution of measureable bacterial count values is given by

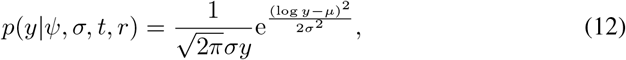

where we set 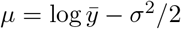 such that the mean error of the candidate models is zero, i.e. 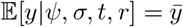.

### 2.6 Data & inference

We estimate the model parameters, (*ψ, σ*): 1. from an *in vitro* static time kill dataset of *N. gonorrhoeae* under the exposure of ciprofloxacin published in [38]; 2. from a synthetically generated dynamic time kill dataset. The *in vitro* dataset contains bacterial density measurements of 12 *N. gonorrhoeae* populations over a time period of 10 h. After an initial growth phase of 4 h, the populations were exposed to different concentrations of ciprofloxacin (see Figure 3A). These concentrations were constant over time. From this dataset, the model parameters are estimated using a maximum likelihood approach as defined in Equation 3. We constrain parameters to realistic orders of magnitudes using upper and lower bounds. The bounds as well as the estimated MLEs are reported in the SI Table A.

**Figure 3:**
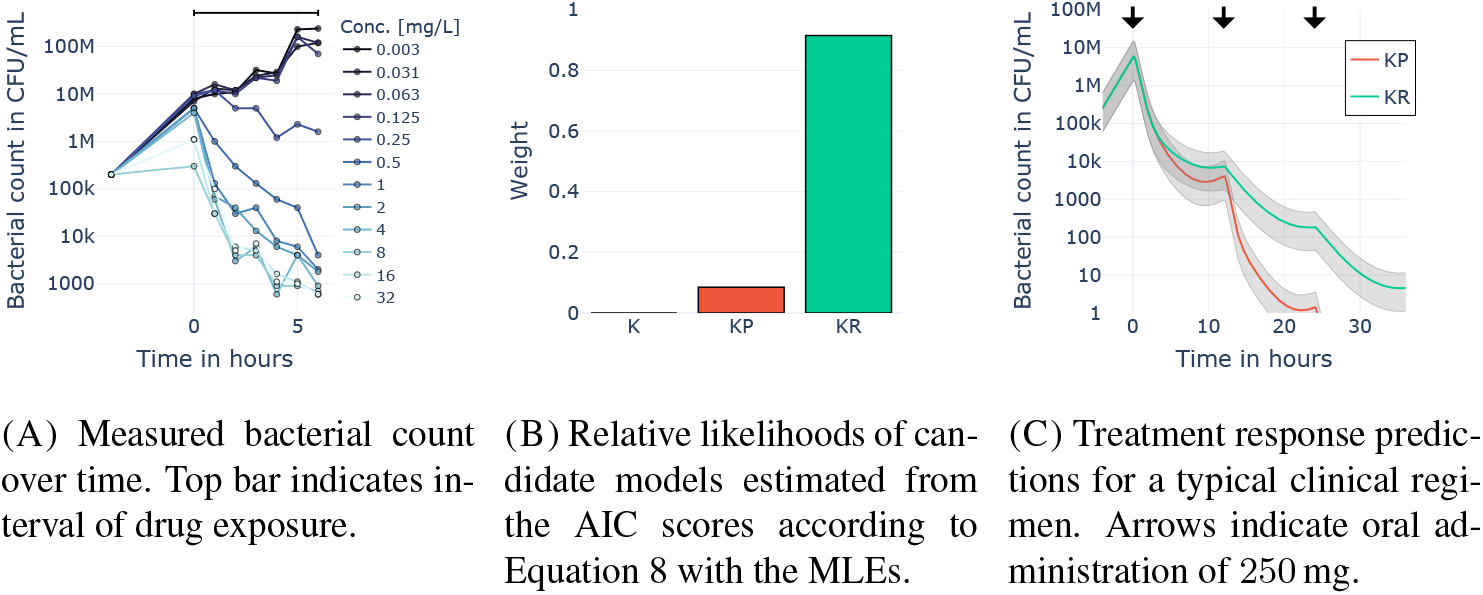
Structural uncertainty after fitting models to *in vitro* static time kill data of *N. gonorrhoeae* under the exposure of ciprofloxacin. (A) shows the data used for the inference, and (B) shows the relative likelihoods of the candidate models. In (C) the treatment response for a clinical dosing regimen is predicted with the two top candidate models. Solid lines indicate the means and shaded areas the 5th to 95th percentiles of the model predictions.

The synthetic dataset is generated using the KR model distribution defined by Equations 10-12, Table 1 and the MLEs derived from the *in vitro* dataset. Measurements are sampled from the distribution hourly for a duration of 30 h for 7 different dosing regimens (see SI Figure A). The dosing regimens were chosen to mirror the dynamics of the *in vitro* experiment for the first 10 h. In particular, the drug is administered after an initial 4 h growth phase for 6 h at infusion rates of different levels. The infusion of the drug is repeated in 15 h intervals. Throughout the experiment, the drug is eliminated from the system at a constant rate (see SI Section B.1 for details). From this dataset, we infer the parameter values 4 times using a Bayesian inference approach as defined in Equation 5. We estimate the parameters: 1. using only the data up to 10 h; 2. using only the data up to 15 h; 3. using only the data up to 20 h; and 4. using only the data up to 30 h. We bound the parameters to realistic orders of magnitudes using uniform prior distributions. The prior distributions as well as the estimated posterior distributions are reported in the SI Tables D-G.

### 2.7 Software & implementation

We implement the mechanistic models in SBML format [39] and simulate the models with the open-source Python package Myokit [40]. The inference is performed using the open-source Python package Pints [41] and its implementations of the CMA-ES and the ACMC algorithms. CMA-ES is used for the estimation of the MLEs from the *in vitro* data, and ACMC is used for the inference of the posterior distributions from the synthesised data. The convergence of the ACMC chains is analysed using the 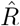 statistic implemented in the open-source Python package ArviZ [42]. Details on the number of warmup iterations, the total number of iterations, the number of chains and the thinning ratio can be found in the SI. The data and models, as well as executable scripts to reproduce the analyses are hosted on GitHub: https://github.com/DavAug/PAM.

## 3 Results

In this section, we report the inference results from the *in vitro* static time kill data and the results from the synthesised dynamic time kill data.

### 3.1 Inference results from in vitro data

The inference results from the *in vitro* static time kill data are illustrated in Figure 3. Figure 3A shows the bacterial count measurements under the exposure of different ciprofloxacin concentrations on a log-scale. Drug concentrations above a threshold appear to induce a decline in the bacterial count throughout the period of exposure. The rate of decline is fastest immediately after the start of the administration and slows down over time. This observation hints that the response dynamics may not be adequately described by the K model, since, in contrast to the KP and KR models, the K model does not provide a mechanism for an adaptation to the drug exposure.

The relative model likelihoods in Figure 3B confirm this intuition and assign negligible weight to the K model, while the KP and KR models receive substantial weights. The weights are estimated according to Equation 8 using the MLEs inferred from the *in vitro* data. The KR model has the highest weight (i.e. the lowest AIC score) and is, thus, chosen for future treatment response predictions by the MS modelling strategy. As a result, MS disregards the considerable structural uncertainty that remains after fitting to the data.

A closer look at the treatment response dynamics of the fitted KP and KR models shows that the population dynamics of persistent and resistant bacteria are qualitatively the same when exposed to constant drug concentration levels (see e.g. first halves of middle and right panels in Figure 4). Only upon re-exposure to sufficiently high drug concentrations do the dynamics of the models start to differ substantially (see e.g. second halves of the bottom panel in Figure 4). Hence, the constant drug concentrations in the *in vitro* study cannot distinguish the KP and KR models as long as the resolution of the observed dynamics is limited by measurement noise. This suggests that the difference in the model weights is not indicative of the true response mechanism, but may in fact be dominated by measurement noise.

**Figure 4:**
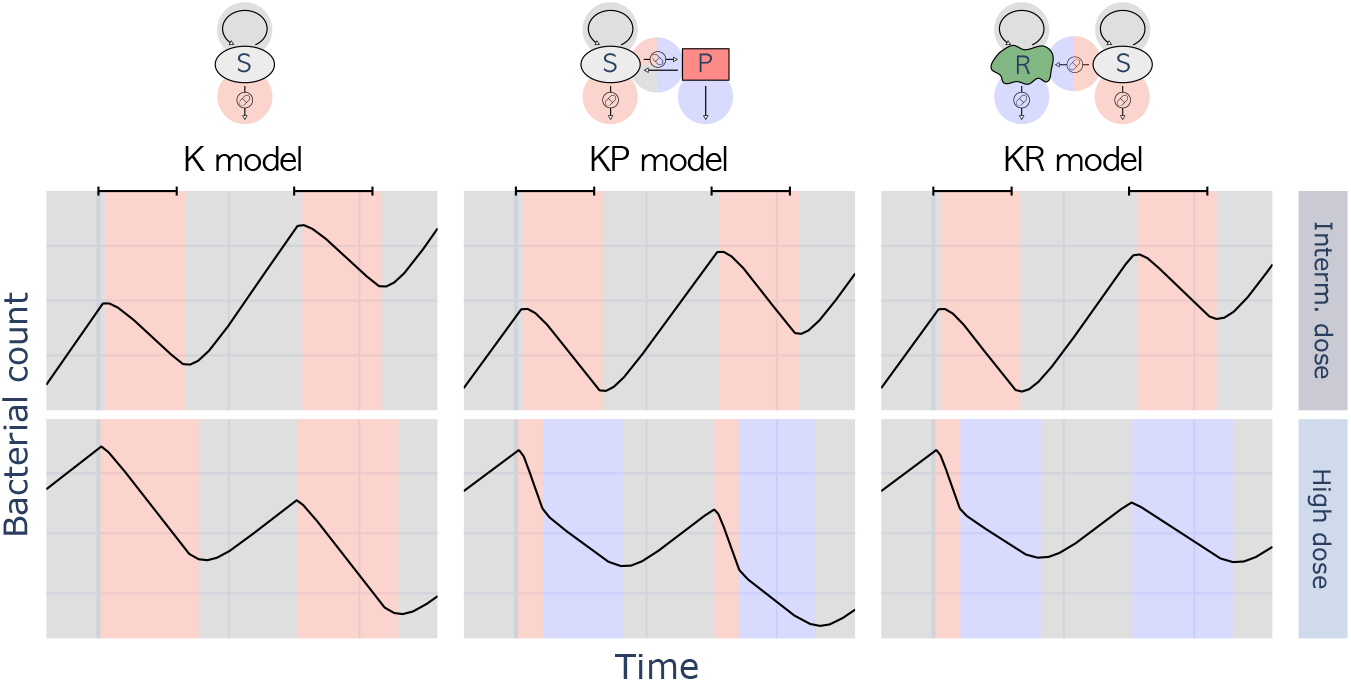
Qualitative treatment response dynamics of the candidate models after fitting to the *in vitro* data in Figure 3A (see SI Section A.4 for details). Distinct dynamical phases are coloured according to the dominant mechanism of the candidate models. Top bars indicate periods of drug infusion at a constant rate.

The consequences of choosing either the KP or the KR model for treatment response prediction are demonstrated for a typical clinical dosing regimen of ciprofloxacin in Figure 3C. Here, the bacterial infection of a virtual patient is treated with a bidaily oral ciprofloxacin dose of 250 mg. The treatment response is predicted with the candidate models and their MLEs. The PK of ciprofloxacin is modelled for an average human (70 kg) according to [43] (see SI Section A.5 for details). Figure 3C shows that the KP and KR models make similar predictions about the treatment response up to the second dose. However, for later times the model predictions start to differ, indicating that neither model is reliable past the second dose. MS disregards this uncertainty by choosing the KR model, regardless of the substantial relative likelihood of the KP model. In contrast, PAM considers both the KP and the KR models, which reflects the remaining structural uncertainty in the treatment response predictions.

### 3.2 Predictive reliability of MS, MAA and PAM

The true response mechanism of the bacteria in the *in vitro* study is unknown which makes it difficult to quantify the significance of the different modelling strategies for the predictive reliability. To overcome this limitation, we synthesise data from a known model and use this to assess the three modelling strategies: MS, MAA and PAM. In particular, we use the KR model and its MLEs from the *in vitro* study to synthethise bacterial count measurements and use the true treatment response as the gold standard (see Section 2.6).

From this dataset, we infer the parameter values for four different dataset durations, with data obtained until 10 h, 15 h, 20 h and 30 h after the start of the experiment. We then predict a typical dose selection endpoint — the dose amount that leads within 24 h to a 1-log (10-fold) reduction of the bacterial population [44] — using the different model selection/averaging strategies (see SI Sections B.2 and B.3 for details). The drug is orally administered twice a day to a virtual patient, similar to the clinical regimen in Figure 3C.

The predicted 1-log reduction doses of the MS, MAA and PAM modelling strategies are illustrated in the top panel of Figure 5. The reliability of the predictions is assessed for the MAA approach based on the proximity to the true 1-log reduction dose, and for the MS and PAM approaches based on the assigned probabilities to the true 1-log reduction dose. The predictions in Figure 5 show that all modelling strategies predict the true 1-log reduction dose reliably when sufficient data are available (i.e. for experiment durations *≥* 15 h). With insufficient data, MAA performs poorly and overestimates the true 1-log reduction dose by almost 100 mg, while MS and PAM still assign the correct dose with a high probability. However, MS does not reliably identify the true dose response mechanism. In fact, MS favours the KP model after both 15 h and 20 h of experimentation. This unreliable model selection aligns with the intuition developed in Section 3.1 that the data cannot distinguish the KP and KR dynamics before the bacteria are exposed to the drug a second time. It also confirms that the differences in the model weights are dominated by measurement noise when the candidate models are close approximations of the observed dose response. After the response of the bacteria to the second infusion is observed, the model likelihoods shift all weight to the correct (KR) model structure.

**Figure 5:**
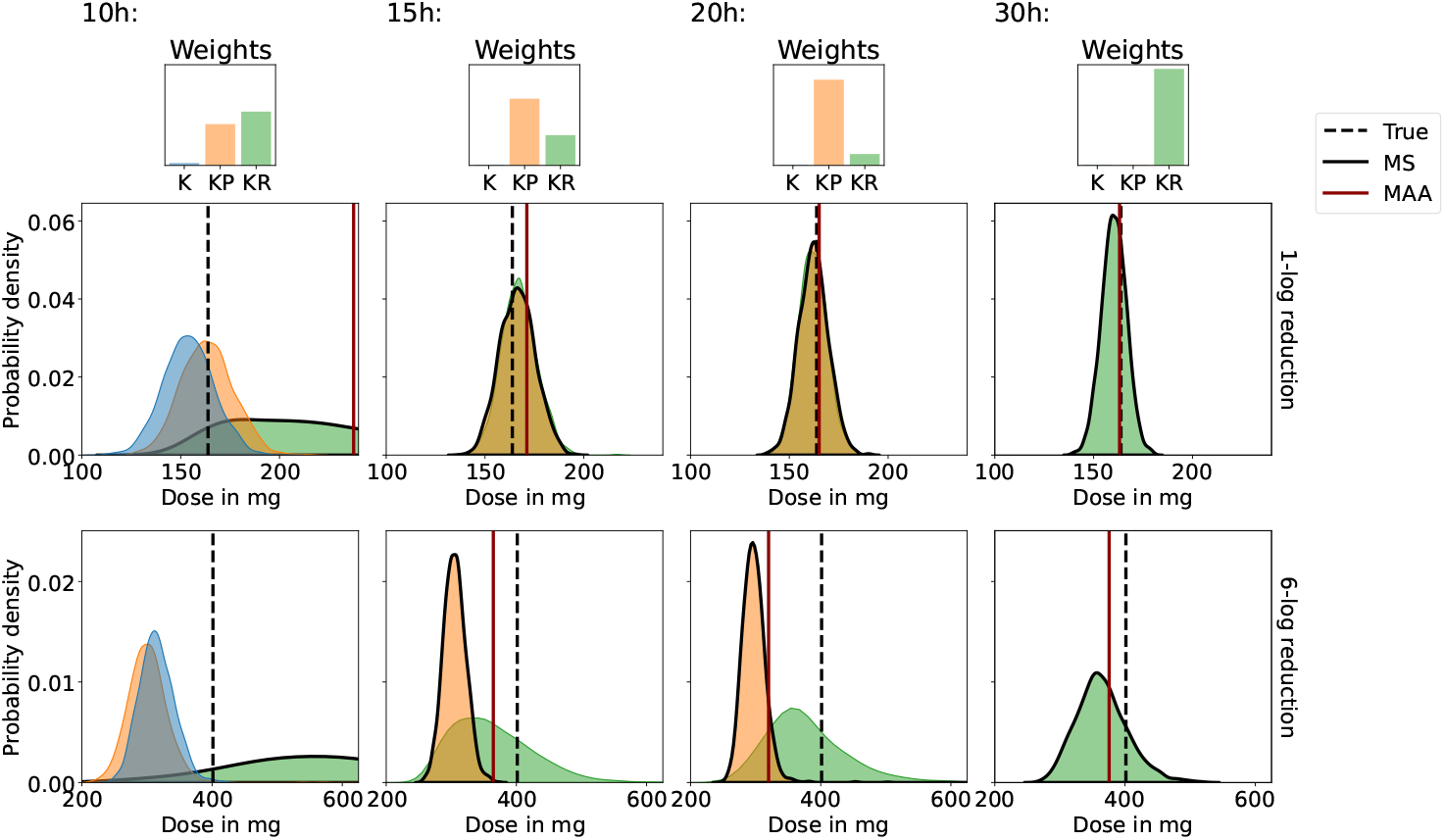
Comparing model predictions for ciprofloxacin treatment of *N. gonorrhoeae* infection. The top panel shows the predictions for the dose that leads within 24 h to a 1-log reduction of the bacterial count. The bottom panel shows the predicted 6-log reduction doses. The columns show the predictions for different durations of the *in silico* experiment: 10 h, 15 h, 20 h and 30 h. The true doses are indicated by dashed lines. PAM’s predicted dose distributions are illustrated by the individual model predictions and the corresponding model weights. Only the distributions from models with a weight above *w*_thresh_ = 10^−3^ are shown. The distributions selected by MS are highlighted with a black edge. MAA’s predicted doses are visualised by solid red lines, except the predicted 6-log reduction dose for 10 h, where the predicted 790 mg is outside the illustrated dose range.

Selecting the wrong candidate model to predict the 1-log reduction dose does not influence the predictions in this instance, because the administered dose is too small to trigger the resistance mechanism of the true treatment response. Thus, the data-generating KR model effectively reduces to a K model, whose dynamics can be accurately described by all candidate models (similar to top panel in Figure 4). However, for dosing regimens that trigger the resistance response of the bacteria, the modelling strategy significantly impacts the reliability of the treatment response predictions (see the bottom panel in Figure 5). Here, we predict the 6-log reduction dose, which requires doses of ciprofloxacin that induce resistant mutations in the bacterial population. Now, MS and MAA only lead to reliable predictions when the data-generating model has been conclusively identified. When insufficient data are available to eliminate the structural uncertainty, both MS and MAA modelling strategies predict false 6-log reduction doses. In contrast, PAM assigns high probability to the correct 1-log and 6-log reduction doses at all times and dataset sizes.

## 4 Discussion

Our results show that the reliable prediction of treatment response dynamics crucially relies on the identification of the correct response mechanisms. This is because the dynamics of biological and pharmacological processes do not trivially extrapolate to the future or unobserved dosing regimens due to their nonlinear nature. This makes PKPD processes challenging to predict unless the true dose response mechanisms are well approximated.

MS attempts to identify the true treatment response mechanism by choosing the model that captures the most information about the observed dynamics. However, as demonstrated in Section 3, it is possible that multiple candidate models closely approximate the observed dynamics, in which case the data leaves structural uncertainty about the candidate models. MS neglects this uncertainty by selecting a single ‘best’ model, which can lead to unreliable model predictions. This is exacerbated by the fact that the differences in the models’ AIC scores can be dominated by measurement noise when multiple models approximate the data well, which leads to an effectively random ranking of candidate models (see e.g. the model weights in Figure 5). We therefore question the appropriateness of MS for treatment response prediction when the data leaves uncertainty about the correct model choice.

MAA attempts to incorporate structural uncertainty about the candidate models by averaging their mean predictions using the estimated model weights, but provides no overall uncertainty quantification. While this strategy can work well, MAA will in general extrapolate poorly. This is because, in contrast to the individual candidate models, the weighted average of the model predictions is no longer founded in a mechanistic understanding of the biological and pharmacological processes. As a result, MAA cannot be expected to predict nonlinear treatment responses well for times or dosing regimens that go beyond the observed treatment response dynamics. However, MAA can lead to good predictive performance when all likely candidate models reduce to qualitatively the same response dynamics for the predicted times and dosing regimens.

PAM remedies the problems of MS and MAA (see e.g. Figure 5). PAM rescales the probability densities of the candidate models relative to their likelihoods. This leaves the predictions of the candidate models unaltered, and therefore preserves the mechanistic justification of the model predictions. In addition, this allows PAM to quantify the overall uncertainty of the predicted treatment response dynamics. When sufficient data are available, PAM reduces to the MS modelling strategy by eliminating unlikely candidate models and assigning all weight to a single dominant model. This makes PAM not only a reliable choice for treatment response predictions, but it may also help to optimise experimental designs.

In this article, we have demonstrated that PAM can be a reliable approach for treatment response predictions, but there are two important limitations of the approach. First, the structural uncertainty, as estimated by the weights in Equation 8, estimates the relative likelihoods of the candidate models under the assumption that at least one candidate model generated the observed treatment response dynamics. This assumption may not be justified for most biological modelling problems where the models only approximate the reality. As a result, the model predictions of PAM are only as reliable as the quality of the candidate models. If all models are poor approximations of the biology, the least poor model is likely assigned with all model weight, ascribing false confidence to the model choice. It is therefore crucial that candidate models are proposed based on pharmacological and biological understanding, and that model fits are visually inspected, e.g. using posterior predictive checks [45], to identify obvious model inadequacies.

Second, PAM’s (as well as MS’s and MAA’s) ability to estimate the relative likelihoods of candidate models is limited by the sensitivity of the AIC scores to measurement noise. As discussed in Section 3, this sensitivity can lead to an arbitrary ranking of candidate models when multiple models capture the observed dose response dynamics. We therefore suggest considering all models with substantial weights equally and disregarding candidate models only below a lower threshold, e.g. *w*_thresh_ = 10^−3^. For more robust estimates of the model weights one may also consider replacing the AIC score by other strategies to estimate the cross entropy, such as cross validation, PSIS-LOO or the WAIC score [46, 47].

In summary, PAM is a modelling framework that reflects the uncertainty of predictions more truthfully than both MS and MAA and may therefore be favoured for drug development, dosing regimen selection and treatment response prediction.

## 5 Study highlights

### What is the current knowledge on the topic?

In PKPD modelling, the data are often insufficient to conclusively determine an optimal model. This uncertainty about the model choice is commonly addressed by choosing the model that is most consistent with the observed data (model selection). More recently, it has also been proposed to average the candidate model predictions based on their relative likelihoods (model averaging).

### What question did this study address?

How reliable are model selection and model averaging for treatment response prediction, and how can we do better?

### What does this study add to our knowledge?

We show that model selection and model averaging can be unreliable for treatment response prediction and dosing regimen recommendation, especially when multiple models are close approximations of the observed treatment response. We demonstrate that probabilistic averaging of models (PAM) generally improves the reliability of predictions.

### How might this change clinical pharmacology or translational science?

PAM makes more reliable predictions of treatment responses, leading to more informed decisions in clinical pharmacology and translational science.

## Supporting information

Supplemental description of methods

## Footnotes

1 Note that **x**, *ψ* and 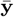 are vector-valued, i.e. **x** ∈ ℝ^*d*^, *ψ* ∈ ℝ^*p*^ and 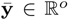, where *d* is the number of dynamical states, *p* is the number of mechanistic model parameters and *o* is the number of observed quantities.

2 The expectation of **y** is only equal to 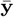 under the common assumption that the model error is on average zero. Other choices are also possible.

3 Note that also *σ* is vector-valued, i.e. *σ* ∈ ℝ^*e*^, where *e* denotes the number of error model parameters.

4 Equation 3 holds under the common assumption that the measurements are independent with respect to the candidate model *p*.

5 Note that even the MLE approach can be expressed as a distribution over model parameters by defining a distribution that is delta-distributed at the MLEs, i.e. 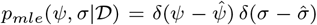. This makes it obvious that the MLE approach neglects parametric uncertainty of the estimates 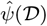 and 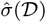.

6 Note that the model parameters *ψ, σ* are, in general, different across candidate models and do not have to have the same dimension. For ease of notation, we will nevertheless refrain from indexing the parameters with *m*.

7 A lognormal error model is equivalent to a Gaussian error on the log-scale 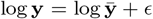, where *ϵ* is a Gaussian random variable with mean 0 and standard deviation *σ*, if the location of the lognormal distribution is chosen to be *µ* = 0.

