## Supplemental description of methods for "Treatment response prediction: Is model selection unreliable?"

### Supplementary information

The raw data, the data derived during the inference, model specifications (SBML format) and executable scripts (Jupyter notebooks) used to produce the results of this article are published on GitHub: https://github.com/DavAug/PAM. This allows readers to investigate all results and the inference pipeline directly in the notebooks. Below, we provide the information contained in the notebooks in a more 'traditional' format.

#### A Study of in vitro static time kill data

##### A.1 Inference of K model

The K model, as defined in Equation 10, Equation 11 and Table 1 in the main document, can be more explicitly written as

$$\frac{\mathrm{d}x_s}{\mathrm{d}t} = \lambda_s x_s - \frac{k_s c}{c + c_{s,50}} x_s \quad \text{and} \quad \bar{y} = x_s, \tag{A}$$

where $x_s$ is the count of the susceptible bacterial population, $\lambda_s$ is the net-growth rate, $k_s$ is the maximal drug-induced death rate, $c$ is the drug concentration and $c_{s,50}$ is the half-maximal effect drug concentration (EC50). The mechanistic model prediction of the bacterial count $\bar{y}$ is equal to $x_s$ for the K model, as the model assumes that there is no heterogeneity in the population (all bacteria are susceptible). The mechanistic model parameters $\psi$, which need to be estimated from the data, are $\psi = (x_{s,0}, \lambda_s, k_s, c_{s,50})$, where $x_{s,0}$ is the initial bacterial count. The error of the K model is modelled with a

*Corresponding author.
†

| Parameter | Unit | Bounds | Estimate | AIC |
|---|---|---|---|---|
| $x_{s,0}$ | CFU/mL | $[0, 10^6]$ | $2.91 \times 10^5$ | |
| $\lambda_s$ | 1/h | $[0, 5]$ | 0.81 | |
| $k_s$ | 1/h | $[0, 5]$ | 2.70 | 2766 |
| $c_{s,50}$ | mg/L | $[0, 10^4]$ | $4.27 \times 10^2$ | |
| $\sigma$ | no units | $[0, 3]$ | 1.40 | |

(1) K model inference results.

| Parameter | Unit | Bounds | Estimate | AIC |
|---|---|---|---|---|
| $x_{s,0}$ | CFU/mL | $[0, 10^6]$ | $2.53 \times 10^5$ | |
| $\lambda_s$ | 1/h | $[0, 5]$ | 0.78 | |
| $\lambda_p$ | 1/h | $[0, 2]$ | 0.27 | |
| $t$ | L/mg/h | $[0, 10^{-3}]$ | $5.21 \times 10^{-5}$ | 2645 |
| $\tilde{t}$ | 1/h | $[0, 5]$ | 0.52 | |
| $k_s$ | L/mg/h | $[0, 10^{-2}]$ | $3.71 \times 10^{-3}$ | |
| $\sigma$ | no units | $[0, 3]$ | 0.72 | |

(2) KP model inference results.

| Parameter | Unit | Bounds | Estimate | AIC |
|---|---|---|---|---|
| $x_{s,0}$ | CFU/mL | $[0, 10^6]$ | $2.52 \times 10^5$ | |
| $\lambda$ | 1/h | $[0, 5]$ | 0.78 | |
| $\mu$ | L/mg/h | $[0, 10^{-4}]$ | $5.04 \times 10^{-5}$ | |
| $k_s$ | L/mg/h | $[0, 10^{-2}]$ | $3.80 \times 10^{-3}$ | 2640 |
| $k_r$ | 1/h | $[0, 5]$ | 1.62 | |
| $c_{r,50}$ | mg/L | $[0, 10^3]$ | $1.91 \times 10^2$ | |
| $\sigma$ | no units | $[0, 3]$ | 0.71 | |

(3) KR model inference results.

TABLE A
Summary of *in vitro* study inference results. (A), (B) and (C) show the inference results of the K model, KP model and KR model, respectively.

lognormal error model (see Equation 12 in the main document), which adds the scale of the error model $\sigma$ as a model parameter.

We estimate the model parameters, $(\psi, \sigma)$, using maximum likelihood estimation with additional constraints on the parameter values. In particular, we define minimal and maximal values for each parameter to limit the search space to realistic orders of magnitudes, see Table A1. For the optimisation we use Pints' implementation of the CMA-ES algorithm [2]. To guarantee robustness of the estimates we sample the initial points for the optimisation uniformly within the bounds and run the optimisation routine 5 times. The estimates with the largest likelihood are selected as MLEs. As documented in the Jupyter notebooks, the estimated parameter values are consistent up to 6 significant digits across the 5 runs. We can therefore assume that the estimates reported in Table A1 can be reliably reproduced with our inference setup. Table A1 also reports the estimated cross entropy of the K model in the form of the AIC score, which we computed according to Equation 6 in the main document using the MLE model, $p_K(y|\hat{\psi}, \hat{\sigma}, t, r)$.

##### A.2 Inference of KP model

The KP model, as defined in Equation 10, Equation 11 and Table 1 in the main document, can be more explicitly written as

$$\frac{\mathrm{d}x_s}{\mathrm{d}t} = \lambda_s x_s - tcx_s + \tilde{t}x_p - k_s c x_s, \quad \frac{\mathrm{d}x_p}{\mathrm{d}t} = -\lambda_p x_p + tcx_s - \tilde{t}x_p \tag{B}$$

and $\bar{y} = x_s + x_p$. $x_s$ and $x_p$ are the counts of the susceptible and the persistent bacterial populations. $\lambda_s$ is the net-growth rate of the susceptible population in the absence of the drug, and $k_s$ is the drug-induced death rate. $\lambda_p$ is the net-death rate of the persistent population, and $t$ and $\tilde{t}$ are the transition rates from the susceptible to the persistent state, and the persistent to the susceptible state, respectively. Note that we modified the drug-induced death of the susceptible bacteria in comparison to Equation A. This is because in the *in vitro* study the data is insufficient to resolve the sigmoidal potency of the drug, when we also allow for the transition into a persistent state. The mechanistic model prediction of the bacterial count $\bar{y}$ is equal to $x_s + x_p$ for the KP model, as the model assumes that the population consists of two subpopulations. The mechanistic model parameters are $\psi = (x_{s,0}, x_{p,0}, \lambda_s, \lambda_p, t, \tilde{t}, k_s)$, where $x_{s,0}$ and $x_{p,0}$ are the initial bacterial counts. The error of the KP model is modelled with a lognormal error model (see Equation 12 in the main document), which adds the scale of the error model $\sigma$ as a model parameter. We estimate the model parameters analogously to Section A.1. However, we fix the initial population size of the persistent bacteria to be zero, $x_{p,0} = 0$. Please find the summary of the inference results in Table A2.

##### A.3 Inference of KR model

The KR model, as defined in Equation 10, Equation 11 and Table 1 in the main document, can be more explicitly written as

$$\frac{\mathrm{d}x_s}{\mathrm{d}t} = \lambda x_s - \mu c x_s - k_s c x_s, \quad \frac{\mathrm{d}x_r}{\mathrm{d}t} = -\lambda x_r + \mu c x_s - \frac{k_r c}{c + c_{r,50}} x_r \tag{C}$$

and $\bar{y} = x_s + x_r$. $x_s$ and $x_r$ are the counts of the susceptible and the resistant bacterial populations, and $\lambda$ is the net-growth rate of the subpopulations in the absence of the drug. $k_s$ is the drug-induced death rate of the susceptible bacteria, and $k_r$ is the maximal drug-induced death rate of the resistant bacteria. $c_{r,50}$ is the half-maximal effect drug concentration (EC50), and $\mu$ is the mutation rate. The mechanistic model prediction of the bacterial count $\bar{y}$ is equal to $x_s + x_r$ for the KR model, as the model assumes that the population consists of two subpopulations. The mechanistic model parameters are $\psi = (x_{s,0}, x_{r,0}, \lambda, \mu, k_s, k_r, c_{r,50})$, where $x_{s,0}$ and $x_{r,0}$ are the initial bacterial counts. Note that, in principle, the resistant bacteria could have a different growth rate than the susceptible bacteria. However, the *in vitro* data does not measure the bacterial growth in the absence of ciprofloxacin after the populations have been exposed to the drug. As a result, the natural net-growth rate and the drug-induced kill rate of the resistant bacteria can only be estimated up to a scaling factor. We therefore make the assumption that the mutations of the resistant bacteria do not affect their growth behaviour in the absence of the drug, which removes this non-identifiability.

| Parameter | Unit | Value |
|---|---|---|
| $a_{d,0}$ | mg | 0 |
| $a_{c,0}$ | mg | 0 |
| $k_a$ | 1/h | 2.7 |
| $k_e$ | 1/h | 0.17 |
| $v_c$ | L | 259 |

TABLE B
Ciprofloxacin PK model parameters for a human with body weight of 70 kg [3].

The error of the KR model is modelled with a lognormal error model (see Equation 12 in the main document), which adds the scale of the error model $\sigma$ as a model parameter. We estimate the model parameters analogously to Section A.1. However, we fix the initial population size of the resistant bacteria to be zero, $x_{r,0} = 0$. Please find the summary of the inference results in Table A3.

##### A.4 Schematic illustration of treatment response dynamics

The qualitative treatment response dynamics of the fitted candidate models in Figure 4 in the main document is generated using the MLEs from the *in vitro* data (see Table A). The administered dosing regimens are the same as the dosing regimens in the synthetic dynamic time kill study for population 5 and 7 in Table C. The treatment response is simulated for a duration of 30 h.

##### A.5 Prediction of treatment response for clinical regimen

In order to predict the treatment of an infection with a clinical regimen of ciprofloxacin, we model the PK of ciprofloxacin according to Sanchez et. al. with a one-compartment PK model with indirect dose administration

$$\frac{\mathrm{d}a_d}{\mathrm{d}t} = -k_a a_d + r(t), \qquad \frac{\mathrm{d}a_c}{\mathrm{d}t} = k_a a_d - k_e a_c \tag{D}$$

and $c = a_c / v_c$ [3]. $a_d$ is the drug amount in the dose compartment, and $a_c$ is the drug amount in the central compartment. $k_a$ is the absorption rate, $k_e$ the elimination rate and $r$ is the time dependent dose rate defined by the dosing regimen. The drug concentration is modelled by $c$ which is related to the drug amount in the central compartment by the effective volume of distribution $v_c$. The dose rate is defined by the dosing regimen (250 mg bidaily oral administration), which we model as bolus doses into the dose compartment. In our implementation, we model bolus doses by drug administrations with a duration of $d = 0.001\,\mathrm{h}$, which translates to dose rates $r_0 = 250/0.001\,\mathrm{mg/h}$ that are administered twice a day for a duration of $0.001\,\mathrm{h}$, i.e. $r(t) = r_0 \sum_{k=0}^{\infty} \theta(12k - t)\, \theta(t - (12k + d))$, where $\theta(x)$ is the Heaviside step function.

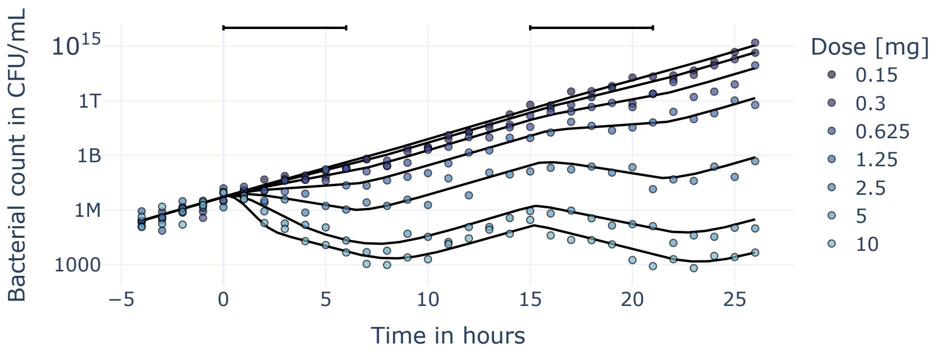


FIGURE A: Synthesised dynamic time kill dataset. Scatter points visualise the measurements of the true treatment response indicated by solid black lines. Top bars indicate periods of drug infusion at a constant rate whose total dose per infusion is indicated in the legend.

### B Study of synthesised dynamic time kill data

#### B.1 Synthesis of data

In this synthetic study, we use the MLE KR model, defined in Equation C, Equation 12 in the main document, and Table A3 to generate measurements of the bacterial count. The PK of the drug is modelled with a one-compartment PK model, where the drug is administered directly into the central compartment

$$\frac{\mathrm{d}a_c}{\mathrm{d}t} = -k_e a_c + r(t) \quad \text{and} \quad c = \frac{a_c}{v_c}. \tag{E}$$

This setup is the model analogue to a hollow fiber experiment where the drug concentrations can be easily modulated [1]. In particular, the drug is cleared from the system at a constant rate $k_e = 1\,\mathrm{h}$, and the volume of the compartment is held at $v_c = 1\,\mathrm{L}$. The drug is administered from $t = 4\,\mathrm{h}$ onwards as infusions of duration $d = 6\,\mathrm{h}$ in $15\,\mathrm{h}$ intervals, i.e.

$$r(t) = r_0 \sum_{k=0}^{\infty} \theta(15k - (t-4))\, \theta(t - 4 - (15k + d)). \tag{F}$$

This dosing regimen is chosen because of its similarity to the constant drug exposure in the *in vitro* study (within $3\,\mathrm{h}$ of the infusion the drug concentration is close to its steady state value $c^* = r_0/k_e/v_c$), but also because it allows for intermediate drug holidays (within $3\,\mathrm{h}$ after the infusion the drug concentration is close to $0$ again.). We measure the bacterial count hourly from $t = 0$ to $t = 30$ for 7 bacterial populations, each administered with a different infusion rate, see Table C.

| Infusion rate | Pop. 1 | Pop. 2 | Pop. 3 | Pop. 4 | Pop. 5 | Pop.6 | Pop. 7 |
|---|---|---|---|---|---|---|---|
| $r_0$ [mg/h] | 0.025 | 0.05 | 0.104 | 0.208 | 0.417 | 0.833 | 1.67 |
| $r_0 \times d$ [mg] | 0.150 | 0.300 | 0.625 | 1.25 | 2.5 | 5 | 10 |

TABLE C
Infusion rates for bacterial populations in *in silico* study. Infusions are administered for a duration of $d = 6\,\mathrm{h}$.

##### B.2 Inference

The parameters of the models are estimated from the synthesised dataset using a Bayesian inference approach (see Equation 5 in the main document). Similar to the *in vitro* study (see Section A), we bound parameters to realistic orders of magnitudes. This is done by formulating uniform priors for the model parameters, see Tables D, E, F and G. The posteriors of all model parameters are estimated using Pints' implementation of the ACMC sampling algorithm [2]. To ensure convergence of the Monte Carlo routine, we initialise 5 chains for each inference problem at random points sampled from the prior distributions. The sampling algorithm is run for 50000 iterations for each chain. The first 10000 iterations are discarded to guarantee that the sampling algorithm has started to sample from high probability regions. To reduce the correlation between samples of nearby iterations, we keep only every 10th iteration. The resulting $4000 \times 5$ samples approximate the posterior distribution of the model parameters. The overall convergence of the chains is evaluated with ArviZ's implementation of the $\hat{R}$ statistic (values close to 1 imply convergence) [4]. The result of the inference process is summarised in Tables D, E, F and G.

The AIC score of the model is computed according to Equation 6 in the main document, where the posterior predictive model is approximated by an average over posterior samples

$$p_m(\mathbf{y}|\mathcal{D}, t, r) \approx \frac{1}{S} \sum_{s=1}^{S} p_m(\mathbf{y}|\psi^s, \sigma^s, t, r). \tag{G}$$

$S$ is the number of posterior samples used to approximate the mean, and $(\psi^s, \sigma^s)$ are the corresponding samples from the posterior distribution.

##### B.3 Prediction of reduction doses

In Figure 5 in the main document, we compare the accuracy of MS, MAA and PAM in predicting the 1-log and 6-log reduction dose of ciprofloxacin. To this end, we determine the true reduction doses by solving

$$\bar{y}(\psi^*, t = 24, r) = \bar{y}_0 \times 10^{-x}. \tag{H}$$

for the dose amount. Here, $\bar{y}(\psi^*, t, r)$ refers to the mechanistic prediction of the data-generating KR model with parameters $\psi^*$ from Section B.1. $\bar{y}_0 = x_{s,0}$ is the initial population size, which we set to $10^6\,\mathrm{CFU/mL}$. $x$ determines the target reduction of the bacterial population, and is therefore either 1 or 6. Similarly to the clinical dosing regimen in Section A.5, the drug is orally administered twice a day. However, the dose amount is no longer fixed at $250\,\mathrm{mg}$, but adjusted to satisfy Equation H. The PK is modelled analogously to Section A.5 with a one compartment PK model with indirect administration. The PK model parameters are again chosen for a human with a body mass of $70\,\mathrm{kg}$, see Table B. To solve Equation H for the dose amount, we minimise the squared distance between the predicted log-reduction and the target log-reduction of the bacterial population using Pints' implementation of the Nelder-Mead optimiser [2]

$$\mathrm{MSE} = \left(x + \log_{10}(\bar{y}/\bar{y}_0)\right)^2 . \tag{I}$$

The true 1-log reduction dose is estimated at $164\,\mathrm{mg}$ and the true 6-log reduction dose is estimated at $401\,\mathrm{mg}$.

**MAA prediction**

The MAA prediction of the reduction doses is estimated analogously to the true reduction doses, see Equation H and Equation I. The main difference is that instead of using the mechanistic model prediction of the data-generating model, we use the MAA prediction according to Equation 7 in the main document. To this end, the expectation of the predictions from the individual candidate models are approximately computed by an empirical mean over the posterior samples

$$\mathbb{E}_m[\mathbf{y}|\mathcal{D}, t, r] \approx \frac{1}{S}\sum_{s=1}^{S} \mathbf{y}^s, \tag{J}$$

where $\mathbf{y}^s$ are samples from the candidate model $p_m(\mathbf{y}|\psi^s, \sigma^s, t, r)$ for the randomly drawn posterior distribution samples $(\psi^s, \sigma^s)$. Each $\mathbf{y}^s$ is drawn for different samples from the posterior distribution. The mean predictions of candidate models are then averaged according to Equation 7 in the main document, where the model likelihoods are estimated from the AIC scores in Tables D, E, F and G to yield the final prediction. This optimisation routine results in point estimates for the reduction doses as illustrated in Figure 5 in the main document.

**MS and PAM prediction**

In order to predict the reduction doses with MS or PAM, we incorporate the uncertainty of the model predictions into the estimation of the reduction doses. This can be done by repeatedly sampling predictions from the candidate models, $p_m(\mathbf{y}|\mathcal{D}, t = 24, r)$, and solving

$$\mathbf{y}(\mathcal{D}, t = 24, r) = \bar{\mathbf{y}}_0 \times 10^{-x} \tag{K}$$

for the dose amount. However, in contrast to $\bar{\mathbf{y}}$ in Equation H, $\mathbf{y}$ is a random variable, so for each sample from $p_m(\mathbf{y}|\mathcal{D}, t = 24, r)$ we get a different estimate of the reduction dose. Repeating this estimation of the reduction dose many times, eventually converges to a stable distribution of reduction doses for each candidate model. This distribution reflects the predictive uncertainty of the models. We repeat this estimation procedure 2000 times for each candidate model.

The MS modelling framework suggests picking the maximum weight model (lowest AIC score) for prediction, while PAM uses all likely models for prediction, see Figure 5 in the main document.

| Parameter | Unit | Prior | Post. mean | Post. std. | $\hat{R}$ | AIC |
|---|---|---|---|---|---|---|
| $x_{s,0}$ | CFU/mL | $U(0, 10^6)$ | $2.5 \times 10^5$ | $0.5 \times 10^5$ | 1.00 | |
| $\lambda_s$ | 1/h | $U(0, 5)$ | 0.83 | 0.04 | 1.00 | |
| $k_s$ | 1/h | $U(0, 5)$ | 3.2 | 0.2 | 1.00 | 2448 |
| $c_{s,50}$ | mg/L | $U(0, 7 \times 10^2)$ | $6.1 \times 10^2$ | $0.6 \times 10^2$ | 1.00 | |
| $\sigma$ | no units | $U(0, 3)$ | 0.82 | 0.07 | 1.00 | |

(1) K model inference results.

| Parameter | Unit | Prior | Post. mean | Post. std. | $\hat{R}$ | AIC |
|---|---|---|---|---|---|---|
| $x_{s,0}$ | CFU/mL | $U(0, 10^6)$ | $2.7 \times 10^5$ | $0.5 \times 10^5$ | 1.00 | |
| $\lambda_s$ | 1/h | $U(0, 5)$ | 0.78 | 0.04 | 1.00 | |
| $\lambda_p$ | 1/h | $U(0, 2)$ | 0.5 | 0.3 | 1.01 | |
| $t$ | L/mg/h | $U(0, 10^{-3})$ | $5 \times 10^{-4}$ | $2 \times 10^{-4}$ | 1.00 | 2442 |
| $\tilde{t}$ | 1/h | $U(0, 5)$ | 0.8 | 0.4 | 1.00 | |
| $k_s$ | L/mg/h | $U(0, 10^{-2})$ | $3.0 \times 10^{-3}$ | $0.3 \times 10^{-3}$ | 1.00 | |
| $\sigma$ | no units | $U(0, 3)$ | 0.76 | 0.06 | 1.00 | |

(2) KP model inference results.

| Parameter | Unit | Prior | Post. mean | Post. std. | $\hat{R}$ | AIC |
|---|---|---|---|---|---|---|
| $x_{s,0}$ | CFU/mL | $U(0, 10^6)$ | $2.8 \times 10^5$ | $0.5 \times 10^5$ | 1.00 | |
| $\lambda$ | 1/h | $U(0, 5)$ | 0.77 | 0.04 | 1.00 | |
| $\mu$ | L/mg/h | $U(0, 10^{-4})$ | $7 \times 10^{-5}$ | $2 \times 10^{-5}$ | 1.00 | |
| $k_s$ | L/mg/h | $U(0, 10^{-2})$ | $3.1 \times 10^{-3}$ | $0.3 \times 10^{-3}$ | 1.00 | 2441 |
| $k_r$ | 1/h | $U(0, 5)$ | 2.0 | 0.3 | 1.00 | |
| $c_{r,50}$ | mg/L | $U(0, 10^3)$ | $6 \times 10^2$ | $2 \times 10^2$ | 1.00 | |
| $\sigma$ | no units | $U(0, 3)$ | 0.76 | 0.06 | 1.00 | |

(3) KR model inference results.

TABLE D
Summary of *in silico* study inference results after 10 h of experiments. (A), (B) and (C) show the inference results of the K model, KP model and KR model, respectively.

| Parameter | Unit | Prior | Post. mean | Post. std. | $\hat{R}$ | AIC |
|---|---|---|---|---|---|---|
| $x_{s,0}$ | CFU/mL | $U(0,10^6)$ | $2.7\times10^5$ | $0.5\times10^5$ | 1.00 | |
| $\lambda_s$ | 1/h | $U(0,5)$ | 0.80 | 0.02 | 1.00 | |
| $k_s$ | 1/h | $U(0,5)$ | 3.1 | 0.1 | 1.00 | 3676 |
| $c_{s,50}$ | mg/L | $U(0,7\times10^2)$ | $6.2\times10^2$ | $0.5\times10^2$ | 1.00 | |
| $\sigma$ | no units | $U(0,3)$ | 0.85 | 0.05 | 1.00 | |

(1) K model inference results.

| Parameter | Unit | Prior | Post. mean | Post. std. | $\hat{R}$ | AIC |
|---|---|---|---|---|---|---|
| $x_{s,0}$ | CFU/mL | $U(0,10^6)$ | $2.9\times10^5$ | $0.4\times10^5$ | 1.00 | |
| $\lambda_s$ | 1/h | $U(0,5)$ | 0.75 | 0.02 | 1.00 | |
| $\lambda_p$ | 1/h | $U(0,2)$ | 0.4 | 0.2 | 1.01 | |
| $t$ | L/mg/h | $U(0,10^{-3})$ | $3\times10^{-4}$ | $2\times10^{-4}$ | 1.00 | 3647 |
| $\tilde{t}$ | 1/h | $U(0,5)$ | 0.7 | 0.2 | 1.01 | |
| $k_s$ | L/mg/h | $U(0,10^{-2})$ | $2.8\times10^{-3}$ | $0.1\times10^{-3}$ | 1.00 | |
| $\sigma$ | no units | $U(0,3)$ | 0.73 | 0.05 | 1.00 | |

(2) KP model inference results.

| Parameter | Unit | Prior | Post. mean | Post. std. | $\hat{R}$ | AIC |
|---|---|---|---|---|---|---|
| $x_{s,0}$ | CFU/mL | $U(0,10^6)$ | $3.0\times10^5$ | $0.4\times10^5$ | 1.00 | |
| $\lambda$ | 1/h | $U(0,5)$ | 0.74 | 0.02 | 1.00 | |
| $\mu$ | L/mg/h | $U(0,10^{-4})$ | $7\times10^{-5}$ | $2\times10^{-5}$ | 1.00 | |
| $k_s$ | L/mg/h | $U(0,10^{-2})$ | $2.8\times10^{-3}$ | $0.1\times10^{-3}$ | 1.00 | 3649 |
| $k_r$ | 1/h | $U(0,5)$ | 1.7 | 0.2 | 1.00 | |
| $c_{r,50}$ | mg/L | $U(0,10^3)$ | $2.3\times10^2$ | $0.9\times10^2$ | 1.00 | |
| $\sigma$ | no units | $U(0,3)$ | 0.74 | 0.05 | 1.00 | |

(3) KR model inference results.

TABLE E
Summary of *in silico* study inference results after $15\,\mathrm{h}$ of experiments. (A), (B) and (C) show the inference results of the K model, KP model and KR model, respectively.

| Parameter | Unit | Prior | Post. mean | Post. std. | $\hat{R}$ | AIC |
|---|---|---|---|---|---|---|
| $x_{s,0}$ | CFU/mL | $U(0, 10^6)$ | $2.8 \times 10^5$ | $0.4 \times 10^5$ | 1.00 | |
| $\lambda_s$ | 1/h | $U(0, 5)$ | 0.81 | 0.02 | 1.00 | |
| $k_s$ | 1/h | $U(0, 5)$ | 3.0 | 0.1 | 1.00 | 5165 |
| $c_{s,50}$ | mg/L | $U(0, 7 \times 10^2)$ | $5.7 \times 10^2$ | $0.5 \times 10^2$ | 1.00 | |
| $\sigma$ | no units | $U(0, 3)$ | 0.88 | 0.05 | 1.00 | |

(1) K model inference results.

| Parameter | Unit | Prior | Post. mean | Post. std. | $\hat{R}$ | AIC |
|---|---|---|---|---|---|---|
| $x_{s,0}$ | CFU/mL | $U(0, 10^6)$ | $2.7 \times 10^5$ | $0.3 \times 10^5$ | 1.00 | |
| $\lambda_s$ | 1/h | $U(0, 5)$ | 0.76 | 0.01 | 1.00 | |
| $\lambda_p$ | 1/h | $U(0, 2)$ | 0.4 | 0.1 | 1.01 | |
| $t$ | L/mg/h | $U(0, 10^{-3})$ | $1.4 \times 10^{-4}$ | $0.8 \times 10^{-4}$ | 1.00 | 5099 |
| $\tilde{t}$ | 1/h | $U(0, 5)$ | 0.5 | 0.2 | 1.00 | |
| $k_s$ | L/mg/h | $U(0, 10^{-2})$ | $3.0 \times 10^{-3}$ | $0.1 \times 10^{-3}$ | 1.00 | |
| $\sigma$ | no units | $U(0, 3)$ | 0.69 | 0.04 | 1.00 | |

(2) KP model inference results.

| Parameter | Unit | Prior | Post. mean | Post. std. | $\hat{R}$ | AIC |
|---|---|---|---|---|---|---|
| $x_{s,0}$ | CFU/mL | $U(0, 10^6)$ | $2.7 \times 10^5$ | $0.3 \times 10^5$ | 1.00 | |
| $\lambda$ | 1/h | $U(0, 5)$ | 0.76 | 0.01 | 1.00 | |
| $\mu$ | L/mg/h | $U(0, 10^{-4})$ | $7 \times 10^{-5}$ | $2 \times 10^{-5}$ | 1.00 | |
| $k_s$ | L/mg/h | $U(0, 10^{-2})$ | $3.0 \times 10^{-3}$ | $0.1 \times 10^{-3}$ | 1.00 | 5103 |
| $k_r$ | 1/h | $U(0, 5)$ | 1.6 | 0.1 | 1.00 | |
| $c_{r,50}$ | mg/L | $U(0, 10^3)$ | $1.7 \times 10^2$ | $0.5 \times 10^2$ | 1.00 | |
| $\sigma$ | no units | $U(0, 3)$ | 0.70 | 0.04 | 1.00 | |

(3) KR model inference results.

TABLE F
Summary of *in silico* study inference results after 20 h of experiments. (A), (B) and (C) show the inference results of the K model, KP model and KR model, respectively.

| Parameter | Unit | Prior | Post. mean | Post. std. | $\hat{R}$ | AIC |
|---|---|---|---|---|---|---|
| $x_{s,0}$ | CFU/mL | $U(0, 10^6)$ | $2.6 \times 10^5$ | $0.4 \times 10^5$ | 1.00 | |
| $\lambda_s$ | 1/h | $U(0, 5)$ | 0.83 | 0.01 | 1.00 | |
| $k_s$ | 1/h | $U(0, 5)$ | 2.63 | 0.06 | 1.00 | 8494 |
| $c_{s,50}$ | mg/L | $U(0, 7 \times 10^2)$ | $4.8 \times 10^2$ | $0.3 \times 10^2$ | 1.00 | |
| $\sigma$ | no units | $U(0, 3)$ | 1.05 | 0.05 | 1.00 | |

(1) K model inference results.

| Parameter | Unit | Prior | Post. mean | Post. std. | $\hat{R}$ | AIC |
|---|---|---|---|---|---|---|
| $x_{s,0}$ | CFU/mL | $U(0, 10^6)$ | $2.3 \times 10^5$ | $0.3 \times 10^5$ | 1.00 | |
| $\lambda_s$ | 1/h | $U(0, 5)$ | 0.780 | 0.008 | 1.00 | |
| $\lambda_p$ | 1/h | $U(0, 2)$ | 0.25 | 0.09 | 1.02 | |
| $t$ | L/mg/h | $U(0, 10^{-3})$ | $2.1 \times 10^{-4}$ | $0.7 \times 10^{-4}$ | 1.00 | 8378 |
| $\tilde{t}$ | 1/h | $U(0, 5)$ | 0.5 | 0.1 | 1.01 | |
| $k_s$ | L/mg/h | $U(0, 10^{-2})$ | $3.02 \times 10^{-3}$ | $0.07 \times 10^{-3}$ | 1.01 | |
| $\sigma$ | no units | $U(0, 3)$ | 0.80 | 0.04 | 1.00 | |

(2) KP model inference results.

| Parameter | Unit | Prior | Post. mean | Post. std. | $\hat{R}$ | AIC |
|---|---|---|---|---|---|---|
| $x_{s,0}$ | CFU/mL | $U(0, 10^6)$ | $2.7 \times 10^5$ | $0.3 \times 10^5$ | 1.00 | |
| $\lambda$ | 1/h | $U(0, 5)$ | 0.767 | 0.007 | 1.00 | |
| $\mu$ | L/mg/h | $U(0, 10^{-4})$ | $7 \times 10^{-5}$ | $2 \times 10^{-5}$ | 1.00 | |
| $k_s$ | L/mg/h | $U(0, 10^{-2})$ | $2.99 \times 10^{-3}$ | $0.06 \times 10^{-3}$ | 1.00 | 8327 |
| $k_r$ | 1/h | $U(0, 5)$ | 1.60 | 0.06 | 1.00 | |
| $c_{r,50}$ | mg/L | $U(0, 10^3)$ | $1.7 \times 10^2$ | $0.3 \times 10^2$ | 1.00 | |
| $\sigma$ | no units | $U(0, 3)$ | 0.71 | 0.03 | 1.00 | |

(3) KR model inference results.

TABLE G
Summary of *in silico* study inference results after 30 h of experiments. (A), (B) and (C) show the inference results of the K model, KP model and KR model, respectively.
